# Techno-functional properties of clover grass protein and the effect of different process technologies

**DOI:** 10.64898/2026.01.27.701969

**Authors:** Narjes Badfar, Simon Gregersen Echers, Charlotte Jacobsen, Betul Yesiltas, Anders Kjær Jørgensen, Tuve Mattson, Peter S. Lübeck, Ankita Mishra, Ana I. Sancho, Katrine Lindholm Bøgh, Mette Lübeck

## Abstract

This study investigated the effects of different downstream processes for protein isolation on the bulk properties and composition of clover grass protein prototypes (CGPs). A clarified clover grass juice, obtained using membrane filtration (MF), underwent precipitation by acid (AP), heat (HP), or acid+heat (AHP), or underwent ultra- and diafiltration to produce a concentrate (DC) as well as subsequent tryptic hydrolysis of DC (DCH). HP had the highest protein content (p<0.05) and was whiter than other CGPs, although it showed lower aqueous solubility. In contrast, DC showed excellent solubility across a broad pH range. CGPs efficiently decreased oil-water interfacial tension (16-13 mN/m) and displayed viscoelastic and solid-like interfacial behavior. CGPs-stabilized emulsions displayed low physical stability with larger droplets despite high absolute ζ-potentials. CGPs were rich in RuBisCO (37-47%) but had varying levels of other proteins. Despite significant protein-level differences, overall protein composition of CGPs was comparable, highlighting that protein state governs bulk functionality more than subtle compositional changes.

**Graphical abstract:** 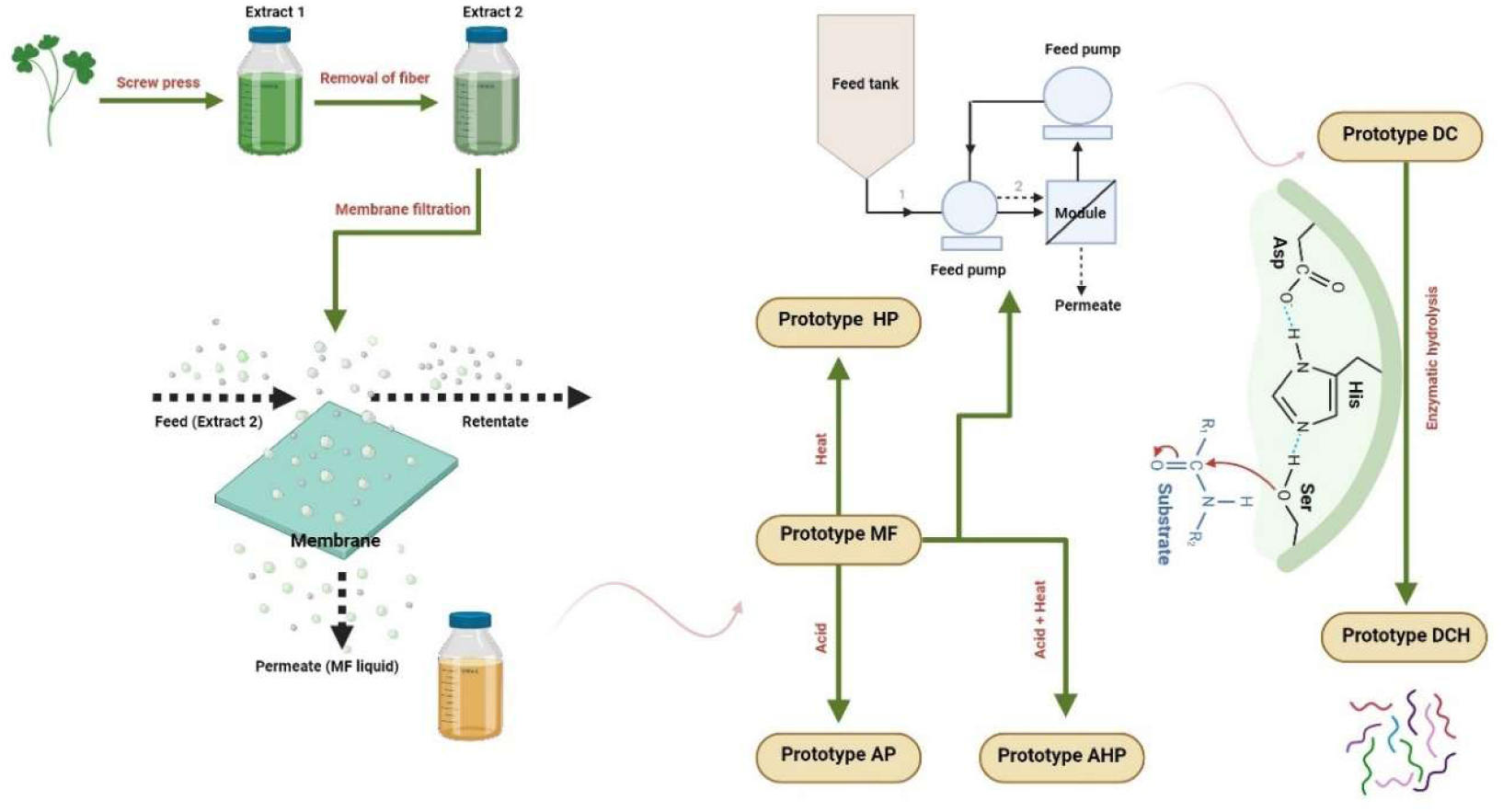

Created with BioRender.com

**Highlights:**

1. The effect of different processes on functional properties of CGPs was explored.
2. Heat treatment increased protein purity and whiteness at the expense of solubility.
3. CGPs efficiently reduced O/W interfacial tension but produced unstable emulsions.
4. CGPs were found rich in RuBisCO (34-47%) using quantitative proteomics.
5. Protein state had larger influence on functionality than protein-level composition.

## 1. Introduction

The world population is growing and is expected to exceed 9.5 billion by 2050 (Henchion et al., 2017). This population increase will necessitate an increased food supply of high nutritional quality, particularly in terms of high-quality protein for human consumption. Given sustainability concerns, the exploration and application of novel protein sources have obtained significant interest and importance (Zhang et al., 2024). Among all sustainable sources, perennial grasses show promise for ensuring a sustainable approach to food supply and reducing the climate impact of the food sector (Pérez-Vila et al., 2024). While perennial grasses and certain green legumes such as clovers and alfalfa are considered sustainable protein sources, they have traditionally been used as animal feed, which is priced lower than protein intended for human consumption (Pearce & Brunke, 2023a; Santamaría-Fernández & Lübeck, 2020). Blends of clovers and grasses (from here on referred to as “clover grass” or “blends”) are widely cultivated in agriculture for animal forages and especially as part of crop rotation schemes in organic farming for soil sanitation and fertilization purposes (Harris & Ratnieks, 2022). The cultivated blends contain various amounts of different species but typically include, but is not limited to, perennial ryegrass (*Lolium perenne*), red clover (*Trifolium pratense*) and white clover (*Trifolium repens*) (Głazowska et al., 2023). The clovers are members of the *Fabaceae* family, characterized by nitrogen-fixation abilities due to bacteria on their roots, which makes them suitable as N supply and in turn helps the production of more protein-rich forages. Therefore, they are known as protein-rich crops (Gallegos Morales et al., 2024).

The nutritional quality of green leaves, in terms of the amino acid composition of the protein, is high and well-suited for human nutrition due to the high content of the essential amino acids, in particular, the higher content of sulphur-containing amino acids, which is low in many plant protein sources on the market, like grains, rice, and peas (Hadidi et al., 2024; Møller et al., 2021; Pearce & Brunke, 2023b). Up to 50% of the soluble leaf proteins are constituted by Ribulose-1,5-bisphosphate carboxylase/oxygenase (RuBisCO), which is the main enzyme involved in photosynthesis (Pérez-Vila et al., 2024). RuBisCO has been shown to have excellent properties as a food ingredient (Di Stefano et al., 2018; Echeverria-Jaramillo et al., 2025; Pérez-Vila, Fenelon, O’Mahony, et al., 2024). Common methods to extract leaf proteins include heat, pH treatment, or lactic acid fermentation; however, they are currently not considered attractive for production of proteins for human consumption, as protein concentrates typically contain bitter tasting compounds, have a relatively low protein content (35-50%), and potentially contain unwanted co-extracted phytochemicals (Santamaría-Fernández & Lübeck, 2020). Heat treatment often results in loss of functionality due to protein denaturation, while lactic acid fermentation causes protein hydrolysis (Nissen et al., 2022). Furthermore, controlling enzymatically driven reactions, such as proteolysis and interactions between activated phytochemicals (e.g. polyphenols) and proteins after initial extraction of green juice, poses a major challenge for implementation of green biorefinery concepts for food protein production at large industrial scale (Møller et al., 2021; Smovzhenko et al., 2025; Tanambell, Danielsen, Devold, et al., 2024; Tanambell, Danielsen, Freund, et al., 2024a). As such, the application of leaf protein in food products remains limited (Hadidi et al., 2023). The effect of different processes for protein isolation on different plant sources’ functional properties has been studied previously. In a comparative study on techno-functionality of pea, chickpea and lentil protein using ultrafiltration and isoelectric precipitation processing techniques, it was reported that protein techno-functionality was directly affected by processing methods but also depended on the plant protein source (Boye et al., 2010). Moreover, the study reported that ultrafiltration preserved native-like structure and charge of the proteins, thus resulting in higher solubility than isoelectric precipitation.

A recent study investigated the application of different green methods for extracting protein from brewer’s spent grains and found that the method and in particular the pH has a significant effect on both yields, purity, and functionality of the protein (Mikkelsen et al., 2025). For RuBisCO-rich green leaf proteins, a mild heat treatment showed a higher solubility and a whey protein isolate-like emulsifying ability and stability at non-acidic pH, compared to non-heat-treated protein isolates (Pérez-Vila, Fenelon, O’Mahony, et al., 2024). Hence, the processing method is a primary driver for obtaining good techno-functional properties of protein. It influences the structure of the protein (aggregation/denaturation, charge, particle size, etc.) and in turn, affects the function of protein. However, no previous studies have systematically investigated how different processing methods affect the composition and techno-functional properties of CGP as a novel food protein from plant sources.

Recently, we developed a gentle, multi-stage membrane separation process, in which a RuBisCO-rich fraction is produced, while the green color from chlorophyll is removed (Gregersen Echers et al., 2026; Mattsson et al., 2025). In this work, soluble proteins from the first stage membrane filtration permeate were recovered by different extraction methods, and the influence on quality of the final protein concentrate and their techno-functionality was investigated. A range of different CGP prototypes, obtained through isoelectric precipitation at slightly acidic conditions, heat precipitation, combined acid and heat precipitation, ultra- and diafiltration, as well as ultra-and diafiltration followed by enzymatic hydrolysis were initially investigated for different techno-functional properties, including solubility, and oil binding capacity (OAC). Next, the protein prototypes, which showed higher solubility and OAC were subjected to further analysis of their emulsifying properties. Emulsification potential was evaluated by investigating physical stability and interfacial properties at the oil-water interface. One of the most broadly examined plant-source proteins is pea protein, as reflected by multiple recent reviews covering its extraction, structure, techno-functional properties, chemical/physical modifications, and diverse applications (Shanthakumar et al., 2022; Shen et al., 2022; Wu et al., 2023). In this study, a commercial pea protein isolate (PPI) was employed as a plant control for comparison with CGP prototypes. In addition, sodium caseinate (Na-Cas) was included as a positive control, since it is widely used in food industries. Prototypes, including the initial and unprocessed green juice, were characterized by quantitative bottom-up proteomics to determine the protein-level composition and how this correlated with the observed techno-functionality. Overall, this study provides novel insight on CGP obtained using different protein isolation methods and links protein state and protein-level composition with functionality. As such, this work can help guide future development of green biorefinery concepts for producing high yield and high quality CGP as a protein-ingredient for foods.

## 2. Materials and methods

### 2.1. Material and sample preparations

Six different clover grass protein prototypes (CGPs) were produced based on different streams of the membrane-based green biorefining of clover grass juice. The clover grass was harvested on the 6^th^ of October 2022 from a field in Jylland, Denmark. The field was planted with a seed blend containing 13% white clover (*Trifolium repens*) and 87% perennial ryegrass (*Lolium perenne*), however red clover (*Trifolium pratense*) was also present in the field. The biomass and initial treatment by maceration and wet fractionation and pre-filtration, followed by crossflow membrane filtration with a 200 nm pore size membrane, as previously described (Mattsson et al., 2025). The permeate from first stage crossflow filtration (MF) was in this work considered the baseline prototype and was subjected to various treatments: acid precipitation (AP), heat precipitation (HP), acid and heat precipitation (AHP), concentrated and purified by ultrafiltration and diafiltration (DC), and finally by tryptic hydrolysis of DC (DCH). DC was produced by crossflow ultra- and diafiltration of the MF fraction over a 10 kDa membrane, as previously described (Gregersen Echers et al., 2026). Compared to our previous study (Mattsson et al., 2025), the MF prototype in this work corresponds to the first stage retentate (S1 Ret) while the DC prototype corresponds to the second stage permeate (S2 Perm) after concentration by diafiltration (DF C). As a reference, the green juice used for initial crossflow filtration (Feed) was included in the proteomics analysis to validate selective retention of undesired and pigment-associated proteins (Gregersen Echers et al., 2026; Mattsson et al., 2025).

All prototypes were freeze-dried in a Telstar LyoQuest freeze dryer (Azbil Co., JP) at -50 °C, 0.500 mBar for approx. 48 h and ground in a Pulverisette table ball mill at 600 RPM (Fritsch, DE). The prototypes were kept at 4 °C in tightly closed containers until further analysis. The pea protein (Emsland group, Emlichheim, Germany) and the sodium caseinate (Arla Foods, Viby J, Denmark) were commercial products.

#### 2.1.1. Preparation of acid or/and heat precipitation process

For HP, the MF baseline fraction was placed in a water bath, preheated to 80°C and kept at that temperature for 1 h. For AP, citric acid (1 M) was added to the MF fraction until a pH of 4 was reached. The solution was then stirred at room temperature (RT) for 1 h. For the combined acid and heat precipitated prototype (AHP), the MF fraction was first subjected to low-pH treatment, followed by the heat treatment, both following the procedures above. After precipitation, all samples were centrifuged at 2000 ×g for 10 min at RT, the supernatant discarded, and the precipitate was freeze-dried to obtain the prototypes AP, HP, and AHP.

#### 2.1.2. Enzymatic hydrolysis of DC concentrate (DCH)

A hydrolysate of DC (DCH) was prepared using the serine protease Formea® Prime (trypsin) 140 kilo microbial trypsin units/gram (KMTU/g) activity (Novonesis (Bagsværd, Denmark) using free-fall pH hydrolysis. The initial pH of the liquid DC was adjusted to 9 with 1 M NaOH and heated to a temperature of 37 °C. Subsequently, the hydrolysis reaction was carried out in an enzyme/substrate ratio (E/S) of 1% (w/w) in a shaking incubator with a constant temperature of 37 °C. The hydrolysis process was continued for 2 h, whereafter enzyme inactivation was accomplished by heating the mixture to 90 °C for 15 min. Afterwards, the mixture was centrifuged at 2000 ×g at 22 °C for 10 min, the supernatant was collected and lyophilized to produce the final DCH.

### 2.2. Physiochemical Characteristics of CGPs

#### 2.2.1. Crude protein analysis by Dumas

The crude protein content was obtained based on the total nitrogen content analyzed by the Dumas method (Rapid MAX N exceed cube N/protein analyzer, Elementar Analysensysteme GmbH, Germany). Measurement performed using 150 mg of the samples. The crude protein (CP) content was determined in all samples by using a factor of 6.25 as reported by previous studies with similar materials (Gregersen Echers et al., 2026; Mattsson et al., 2025) (n=3).

#### 2.2.2. Total amino acid composition using LC-MS

The amino acid composition was determined by HPLC-MS, following hydrolysis and derivatization using EZ:faast amino acid kit (Phenomenex, Torrance, CA, USA) as reported by Jafarpour et al. (2020). Briefly, CGP prototypes were hydrolyzed with 6 M HCl for 1 h at 110°C in a microwave sample preparation system (Multiwave 3000, Anton Paar, Graz, Austria). The subsequent neutralized samples were purified by a solid-phase extraction sorbent tip and derivatization was performed following the injection of sample aliquots into an Agilent HPLC 1100 instrument (Santa Clara, CA, USA) coupled to an Agilent ion trap mass spectrometer. The amino acids were identified by comparing retention time and mass spectra of an external standard mixture. Calibration curves were prepared and analyzed by HPLC-MS for quantification. All analyses were carried out in triplicate samples (with two analytical replicates; n=3×2).

#### 2.2.3. Color assessment

The color assessment of the lyophilized CGP prototypes was conducted using the Hunter Lab Miniscan XE colorimeter (Reston, Virginia, USA). The CIE L*a*b* color parameters were employed for analysis: L*; representing lightness from black (0) to white (100); a*; indicating the degree of redness from green (-120/ negative values) to red (+120 / positive values); and b*; representing the degree of yellowness going from blue (-120 / negative values) to yellow (+120 / positive values) (Badfar et al., 2022). The whiteness value of each prototype was determined as follows:

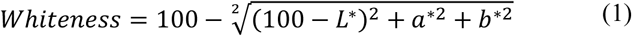

### 2.3. Functional properties

#### 2.3.1. Relative Solubility

To assess the relative solubility of CGP prototypes, 100 mg of powder was dispersed in 10 mL of 0.1 M sodium phosphate buffer (pH 2, 4, 6, 7, 8, 10, and 12). The dispersion was thoroughly mixed by stirring for 10 s, after which the solution was left at RT and shaken at 80 rpm for 30 min. Subsequently, the solutions were centrifuged at 7500 ×g for 15 min. The protein/peptide content in the soluble fraction (supernatant) was determined using the Dumas method as described above. The solubility was calculated as follows (Badfar et al., 2025):

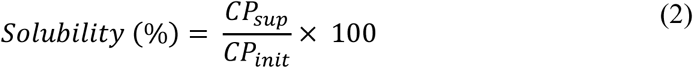

Where CP_sup_ is crude protein content (by Dumas) in the supernatant and CP_init_ is total amount of added crude protein.

Electrophoretic characterization of protein solubility was performed by SDS-PAGE analysis under reducing conditions, as previously described (Danner Aakjaer Pedersen et al., 2025; Mattsson et al., 2025). To ensure comparability, each lane was loaded with an equal volume (5 µL, corresponding to 50 µg protein (by Dumas) if fully solubilized).

#### 2.3.2. Oil Absorption Capacity

Oil absorption capacity (OAC) was determined following the method of Jafarpour and colleague (Jafarpour et al., 2020) with some modifications. Briefly, OAC was determined by dispersing 100 mg of CGPs in 1000 µL of rapeseed oil for 30 s. The resulting mixtures were incubated at RT for 30 min. The samples were centrifuged at 13600 ×g for 10 min. Free oil was decanted and the OAC was calculated as,

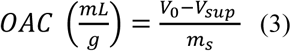

Where V_0_ is the initial oil volume (mL), V_sup_ is the volume of unabsorbed oil in the supernatant after centrifugation/drain (mL), and m_s_ is the mass of the protein content in prototype (g).

### 2.4. Interfacial properties at the water/oil interface

#### 2.4.1 Interfacial tension and dilatational rheology

Interfacial tension and dilatational rheology tests were carried out using a drop tensiometer (OCA20, DataPhysics Instruments, Filderstadt, Baden-Württemberg, Germany) equipped with an oscillating drop generator (ODG20, Dataphysics). A rising droplet of protein solution (35 μL) with a protein concentration of 0.1% (w/v) was formed at the tip of a needle (d=1.83 mm) in an MCT (medium chain triglyceride)-oil-phase within a glass cuvette. The interfacial tension was performed for at least 30 min at RT while keeping the droplet volume constant to reach equilibrium. To determine interfacial tension (γ) based on the Young-Laplace equation, a CCD camera digitized the boundaries of taken droplet images. Regarding performing the dilatational rheology test, the linear viscoelastic regime was determined after a 3 h equilibration. The deformation amplitude was 1.00%, 2.75%, 4.50%, 6.25%, and 8.00% interfacial area at a frequency of 0.01 Hz, where each oscillation was subjected to 5 cycles, and the interfacial film recovered for the 90 s between each step. The frequency sweeps were carried out after the amplitude sweeps, where the frequency varied from 0.01 to 0.10 Hz at an amplitude of 4.00%.

The surface dilatational elastic (E’) and viscous (E’’) moduli were determined from the measured dynamic interfacial tension response, taking the intensity and phase of the first harmonic after Fourier transformation of the γ signal using the equations below (Lucassen and Barnes 1976):

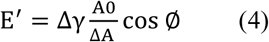

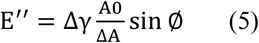

Where Δγ is interfacial tension variation, ΔA is amplitudes of periodic interfacial area, A_0_ represents the unperturbed interfacial area, and ∅ is the phase angle.

### 2.5. Emulsifying properties

#### 2.5.1. Emulsion Preparation

Emulsions were prepared using 0.4 wt% protein of CGPs prototypes, sodium caseinate or pea protein powder in acetate-imidazole buffer (10 mM, pH 7), with subsequent shaken (130 rpm) in a water bath shaker at 50 °C for 2 h, whereafter samples were hydrated overnight at 130 rpm in darkness at RT. The next day, 5% rapeseed oil was added into the aqueous phase and pre-homogenized at first with a handheld ultraturax (Polytron, PT1200E, Littau, Switzerland, 18000 rpm, 30 s) and secondary using a sonicator (Microson XL2000, probe P1) at 75% amplitude (180 μm) on ice for 30 s in two stages with 1 min pause break. The emulsions were stored for 8 days at RT in darkness (n=4).

#### 2.5.2. Zeta Potential, droplet sizes and creaming index

The zeta potential of the emulsions was measured on Day 1 and 8 of storage by ZETASIZER NANO ZS (Malvern instruments Ltd., Worcestershire, UK) with a DTS1070 cell at 25 °C. Ten μL emulsion was diluted in 5 mL Imidazole buffer (10 mM, pH7). The zeta potential range was adjusted to −100 to +50 mV, and the samples were measured with 100 runs. Measurements were carried out in triplicate. The distribution of the droplet size of the CGPs prototypes, pea protein, or sodium caseinate emulsions was recorded by laser diffraction in a MASTERSIZER 2000 (Malvern Instruments, Ltd., Worcestershire, UK) on Day 1, 3, and 8 during storage in the dark at RT. The volume and surface mean diameter (D4,3 and D3,2, respectively) measurements were done in triplicate. The creaming index (CI) was recorded during storage by observing the appearance of the emulsions and calculated as:

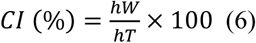

Where hW is the height of the clear water and hT is the total height of the emulsion when stored in a 5 mL measuring tube.

### 2.6. Proteomics analysis by LC-MS/MS

The six prototypes and the initial feed stream for membrane filtration (Feed) were investigated by LC-MS/MS-based bottom-up proteomics, as previously described (Gregersen Echers et al., 2026). Briefly, samples were prepared using a combination of the iST kit for plant tissue (PreOmics, Germany) and focused ultrasonication. A sample mass/volume corresponding to ∼40 µg protein (based on CP by Dumas) was mixed with Lyse buffer to a final volume of 50 µL. Suspensions were heated to 95 °C for 5 min in a Thermomixer (Eppendorf, Germany) at 1000 rpm. Next, the samples (including potential solids) were transferred to 50 µL AFA tubes (Covaris) and subjected to an extraction cycle (peak incident power of 75 W, a duty factor of 10%, 200 cycles per burst, and 180 s per cycle at 6 °C). Subsequently, the liquid phase was transferred to new Protein LoBind tubes (Eppendorf) and heated to 95 °C for 5 min in the thermomixer. After extraction, reduction, and alkylation, proteins were mixed with resuspended Trypsin/LysC and subjected to digestion for 3 h at 37 °C and 500 rpm in the thermomixer. The digest was transferred to iST cartridges where they were subjected to three washing steps before double elution. Lastly, the desalted digest was vacuum dried in a SpeedVac (Thermo-Fisher Scientific, Waltham, MA, USA) and resuspended in LC load buffer.

The analysis was performed on an EASY nLC-1200 ultra-high-performance liquid chromatography system (Thermo-Fisher Scientific) with an ESI ion source coupled to a Q Exactive HF tandem mass spectrometer (Thermo-Fisher Scientific). Approximately 1 μg sample was loaded on a PEPMAP (Thermo-Fisher Scientific) trap column (75 μm x 2 cm, C18, 3 μm, 100 Å) and subsequently separated on a reversed-phase PEPMAP (Thermo-Fisher Scientific) analytical column (75 μm x 50 cm, C18, 2 μm, 100 Å). Peptide separation was accomplished using a step-wise gradient from 5% to 100% buffer B (80% acetonitrile, 0.1% formic acid (VWR, Søborg, Denmark)) in buffer A (0.1% formic acid (Fisher Scientific, Roskilde, Denmark) over 60 min. Data acquisition was performed as full MS/ddMS2 Top20 DDA in positive mode, as previously described (Gregersen Echers et al., 2026).

#### 2.6.1. Analysis of LC-MS/MS data

LC-MS/MS data was analyzed in MaxQuant v2.2.0. 0 (Tyanova et al., 2016). As the initial green juice feed stream was produced from a biomass originating from a field comprised by white clover and perennial ryegrass, with additional red clover observed in the field, all UniProt entries for *L. perenne* (taxid: 4522, 825 entries), *T. repens* (taxid: 3899, 477 entries), and *T. pratense* (UP000236291, 60,146 entries) were used as protein databases. *L. perenne* and *T. repens*, the main constituents, do not have reference proteomes and hence, the reference proteomes for the related *Brachypodium distachyon* (purple false brome, UP000008810, 44,786 entries) and *Trifolium subterraneum* (subterranean clover, UP000242715, 36,725 entries) were used also included in the protein database. Furthermore, as fescues are often found in Danish clover grass fields, all entries from the *Festuca* genus (taxid: 4605, 1,498 entries) were also included. All protein lists were downloaded from Uniprot on December 2nd 2024 (“UniProt: The Universal Protein Knowledgebase,” 2017). Protein identification was performed using standard settings in MaxQuant, as previously described (Gregersen Echers et al., 2026). Protein quantification was done using both the built-in label-free method, MaxLFQ (Cox et al., 2014), as well as intensity-based absolute quantification, iBAQ (Schwanhäusser et al., 2011). The mass spectrometry proteomics data have been deposited into the ProteomeXchange Consortium via the PRIDE (Perez-Riverol et al., 2022) partner repository with the dataset identifier PXD070032 and DOI 10.6019/PXD070032.

#### 2.6.2. Downstream analysis of MaxLFQ data

Quantification by MaxLFQ was analyzed using Mass Dynamics 3.0 (Quaglieri et al., 2022). Missing values were imputed using “missing not at random (MNAR)” with a mean position factor of 1.8 and a standard deviation factor of 0.3. Differential proteins across the entire dataset were identified using the built-in analysis of variance (ANOVA) analysis for heatmap visualization applying row-based, Z-score standardization and protein-level clustering with a Euclidean distance of 6. Pair-wise comparisons were performed based on sample triplicates and visualized in volcano plots. Proteins were considered significantly differential if the adjusted p-value < 0.05 and fold change larger than two (log2FC > 1).

#### 2.6.3. Downstream analysis of iBAQ data

Prior to any computation, the list of 2018 identified proteins (Table S1) was filtered for contaminants (10) and false positive identifications (28). Next, proteins were filtered based on reproducible quantification, requiring that a protein must be reproducibly quantified (i.e. intensity > 0) in at least two replicates of at least one sample. This reduced the list of reproducible proteins to 1740. Overlap of reproducible IDs between streams/prototypes was visualized in an UpSet plot using HiPlot (https://hiplot.cn/basic/upset-plot).

The mean iBAQ intensities between sample replicates were then used to compute the relative, molar protein distribution by dividing the iBAQ for each protein group with the sum of iBAQs for all identified proteins within the sample. The dataset was delimited by including an abundance-based threshold of riBAQ > 0.5% in any sample to only focus on the abundant proteins. This ultimately reduced the initial list of 2018 protein identifications to 63 highly abundant proteins (Table S2).

According to the method described in (Gregersen Echers et al., 2026), the total riBAQ for the following families/groups across the full dataset: RuBisCO (rbc), Chlorophyll a-b binding protein (CBP), Photosystem I & II proteins (PI/II), and non-specific lipid transfer proteins (nsLTPs) was also computed (Table S2). While rbc is the primary target protein, CBP and PI/II were previously shown to co-occur with the green color and grassy sensory attributes, and nsLTPs were highlighted as a food allergen of potential concern (Gregersen Echers et al., 2026).

### 2.7. Statistical analysis

All interfacial tension and rheology experiments were carried out at least twice as independent replicates. ANOVA was performed using the statistical software SPSS Version 28.0.1.1. For significant ANOVA outcomes, means were compared pairwise (p < 0.05) using the Tukey test. From the oscillation experiments, the middle 3 cycles per amplitude were used for the analysis of the rheological moduli. For comparative analysis of protein family/group abundance, statistical analysis was performed as ordinary one-way ANOVA using GraphPad Prism (v.10.0.2, build 232). Comparisons of means were performed with Tukey at 95% confidence intervals assuming Gaussian distribution of residuals and equal standard deviations between groups of unmatched replicates.

## 3. Results and discussions

### 3.1. Proximate protein content and total amino acid composition of CGPs

Based on proximate analysis using Dumas (Table 1), heat precipitation (HP) resulted in the highest protein content (71.58%) and was significantly higher than all other prototypes (p<0.05). The first stage permeate (MF) prototype had significantly (p<0.05) lower protein than remaining prototypes. Heat treatment can aggregate and denature a wide range of proteins and peptides (Van de Vondel et al., 2021); thus, a higher amount of protein precipitated with heating treatment was acquired compared to other methods. In contrast, the first-stage membrane filtration removes residual cell debris, fibers, and chloroplasts, but the resulting permeate and subsequent freeze-dried powder may contain other substances in addition to protein, such as carbohydrates, phenolic compounds and, more importantly, a high level of buffer salts. Therefore, the protein percentage in the resulting baseline prototype powder (MF) is not as high as in the other prototypes which are further treated after initial membrane filtration. Subsequent ultra- and diafiltration concentration (DC), potentially followed by enzymatic hydrolysis (DCH) effectively removed other substances and increased protein content. Conversely, the combination of acidic and heating methods leads to a substantial reduction in the protein content compared to using heat (HP) or acid (AP) precipitation alone, based on Dumas. This discrepancy between protein content obtained by the combination of heat and acid precipitation and individual heat or acid precipitation can be explained by the effects of the other substances that may be formed by heat and acid treatment and nitrogen content measurement by Dumas. In some cases, the inaccuracies were connected to indirect measurements. Nitrogen determination and subsequent conversion to protein, or interference from other chemical substances can affect the results from protein content measurements. Therefore, amino acid analysis is the only method for measuring protein content where interfering substances do not disturb the results (Mæhre et al., 2018). As Dumas relies on standardized conversion of nitrogen to protein that is highly biomass and sample dependent (Gregersen Echers et al., 2026; Mariotti et al., 2008; Yeoh & Wee, 1994), the amino acid composition of all prototypes was determined by LC-MS.

**Table 1.**
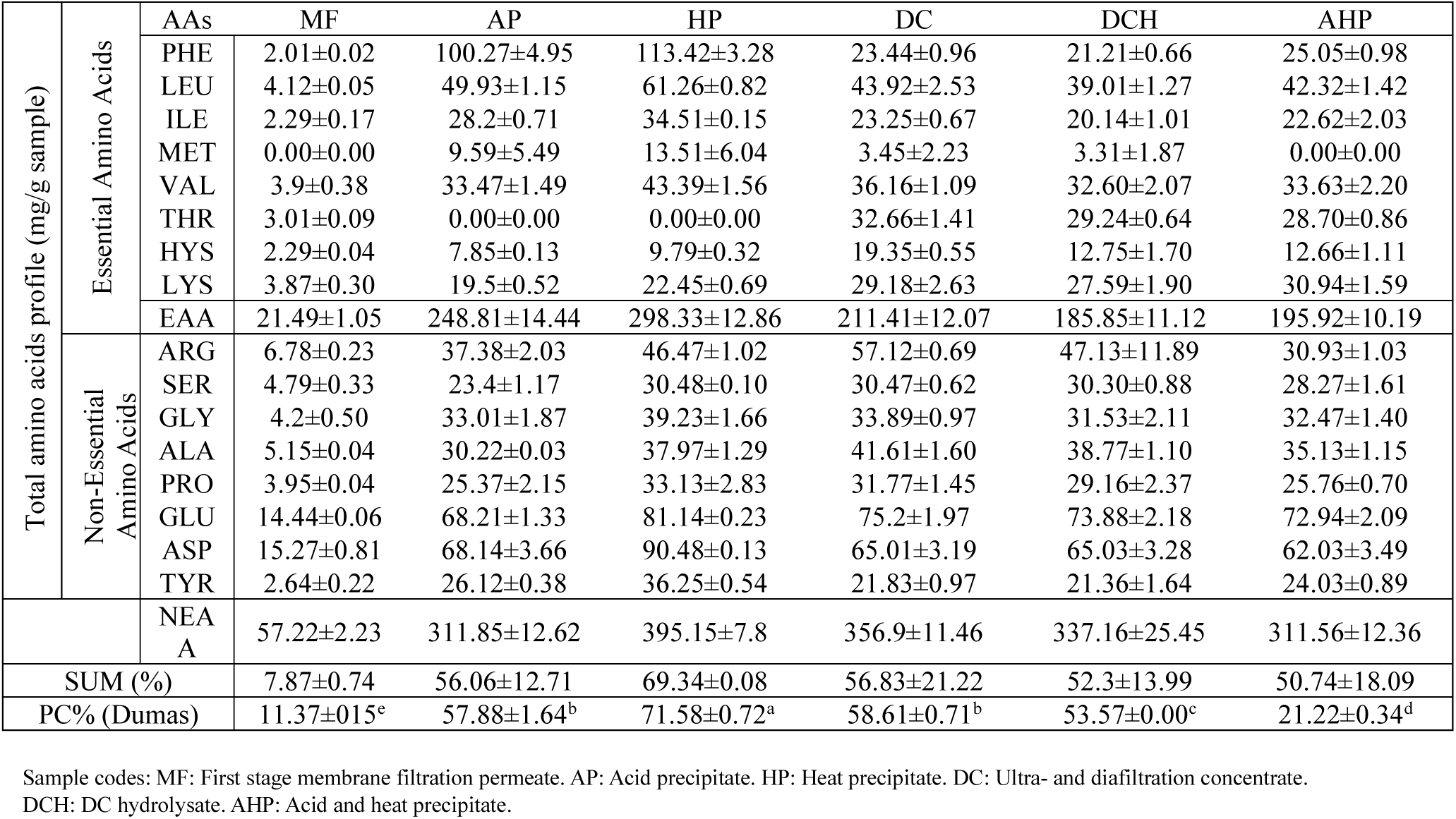
Amino acid composition (mg/g sample) and proximate protein content (% by Dumas N*6.25) of the different clover grass protein (CGP) prototypes investigated.

Results showed that different extraction methods affected the amino acid composition of CGPs differently (Table 1). The amino acid composition results were in line with proximate protein content measured by the Dumas method, except for the AHP prototype. Although the protein content of the AHP prototype was recorded as 21.22% by the Dumas method, the total amino acid content in the same prototype was determined to be 50.74%. This discrepancy between protein content and total amino acid content indicates that the Dumas method may not be accurate for determining protein content when a combination of two methods (AP+HP) have been used to prepare the protein extract (AHP). Amino acid profiles of food proteins are important as these contain both the essential and non-essential amino acids which are crucial for a high-quality diet (Mohanty et al., 2014). Therefore, if the amino acid composition provides especially essential amino acids in appropriate amounts, that protein can be considered as a complete protein (Pérez-Vila et al., 2024). All CGPs obtained from different processing techniques in this study met the needs for essential amino acids (FAO, 2011). However, the level of sulfuric amino acids in all the prototypes was low, as legume proteins generally contain lower levels of sulfur-containing amino acids (Met and Cys) and they are typically considered as limiting amino acid in legume proteins (Huamaní-Perales et al., 2024). According to the results, aspartic acid (Asp) and glutamic acid (Glu) are the most abundant amino acids in CGPs. The same result has previously been reported for white and red clover grass extracts as feed protein (Damborg et al., 2020). However, phenylalanine (Phe) showed the highest value of amino acid content in AP and HP prototypes. Several previous studies investigated amino acid composition of RuBisCO-rich protein isolates/concentrates and reported a similar amino acid profiles from various green leaves (Damborg et al., 2020; Di Stefano et al., 2018; Pérez-Vila, Fenelon, Hennessy, et al., 2024; Zengin et al., 2012). Moreover, (Tanambell, Danielsen, Devold, et al., 2024) confirmed that even under the different extraction and processing techniques, RuBisCO proteins from alfalfa demonstrated high digestibility. Therefore, these findings strengthen and support the potential of green leaves as a well-balanced and nutritious protein source.

### 3.2. Color assessment

To determine the impact of different extraction methods on the color of each prototype, color assessment was carried out using the CIE LAB model (Table 2). The HP prototype showed the darkest (brownish) color compared to the other prototypes with the highest b* and lowest L* and whiteness values. This may be due to heat-induced browning reactions or oxidation of residual phenolic compounds that occurred during the heat precipitation process (Pérez-Vila et al., 2024). The AP prototype, obtained by acid precipitation, was lighter than the HP prototype but darker than the rest of the prototypes (p<0.05). In other words, the two harsh extraction methods (heat or acid precipitation) resulted in a darker prototype powder than the other prototype powder. Interestingly, the combination of these two methods (heat and acid precipitation) resulted in lighter and whiter prototype powder, indicating that some of the proteins, which were attached to the pigments or the other substances remained soluble in the aqueous phase and were not precipitated by application of heating and acidic conditions together or that they were not formed under these conditions. This can be confirmed by total amino acid composition as well, where AHP showed a lower value of total amino acid (50.74%) than HP (69.34%) and AP (56.06%), individually. While the initial membrane filtration process removed some impurities, the MF prototype still contained a large proportion of non-protein content (7.87% total AA content). Following the AHP prototype, the MF prototype was the second lightest and whitest prototype powder among all the prototypes, as it did not undergo additional processing steps and still retained a high content of added buffer salts. Moreover, DC and DCH prototypes exhibited higher L* and whiteness values compared to the HP and AP prototypes, suggesting a lower potential for reactions during these processes. Tanambell, Danielsen, Freund, et al., (2024b) investigated ultrafiltered alfalfa juice and reported color parameters for feed, retentates and permeates. Altering the membrane cut-off from 100 to 300 kDa shifted the color, as a more yellow color was obtained in the100 kDa retentate, suggesting that this is linked to different partitioning of flavonoids. In a study carried out by Mirón-Mérida et al., (2024) on duckweed, it was reported that ultrasound extraction made the extracts darker (lower L*) and more yellow/red (higher a, b*) than the control sample, showing that processing method affects color parameters in a RuBisCO-rich leafy protein. Additionally, Hansen et al., (2022), monitored an alfalfa white protein process and showed changes in L*, a* and b* color parameters profile across mechanical pressing and reported that by increasing pressing steps, the amount of green color increased, suggesting an increase in extraction of the green chlorophyll to the alfalfa protein concentrate.

**Table 2.**
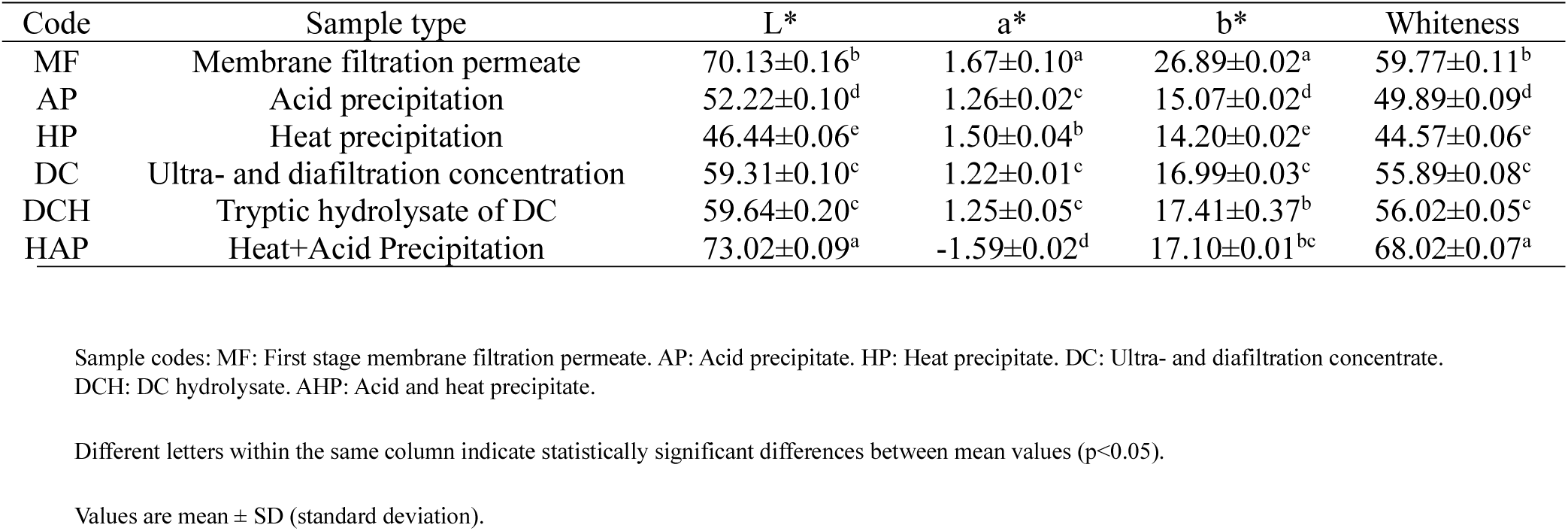
Color parameter assessment of lyophilized clover grass protein prototypes (CGPs).

### 3.3. Functional properties of CGPs

#### 3.3.1. Relative Solubility

Solubility is a crucial functional property in the food industry, directly affecting other properties such as emulsifying, foaming, and gelling abilities (Hayati Zeidanloo et al., 2019). It has been reported that RuBisCO protein has a high relative solubility in water. However, its solubility depends on different parameters such as the isolation process, pH, and mineral composition (Tan et al., 2022). Soluble proteins ensure homogeneous dispersibility in colloidal systems and can enhance interfacial properties (Tarahi & Ahmed, 2023). Consequently, the solubility of CGPs across a pH range of 2-12 was determined (Fig. 1A). According to the results, additional processing of the first stage membrane permeate (MF) leads to varying solubility patterns among different samples. The highest solubility for the MF was observed at pH 7. The highest solubility value for the AP prototype was obtained at pH 8 while solubility was substantially lower at other pH values ranging from 6.4% to 21.59%. Lamsal et al., (2007), reported over 90% solubility of alfalfa leaf proteins under the acid precipitation at the pH 7 before freeze-drying step. Heat treatment resulted in very low solubility across all pH levels, as is expected upon heat-induced denaturation and aggregation of the clover grass proteins (Runyon et al., 2015). A similar pattern was observed with the AHP treatment, suggesting denaturation and aggregation due to heating damage. Conversely, ultra- and diafiltration (DC) and subsequent enzymatic hydrolysis (DCH) demonstrated higher solubility than the rest of samples at pH 8, with solubility values of 77.5% and 89.8%, respectively. Interestingly, DC maintained high solubility even under highly acidic (pH 2, 38%) and highly basic (pH 12, 57%) conditions. High value of solubility at highly acidic and basic pHs for grass pea protein (*Lathyrussativus L.*) isolated by diafiltration was reported by (Hayati Zeidanloo et al., 2019). While various parameters in the hydrolysis process, such as the type of applied proteases, duration, etc. (Vogelsang-O’dwyer et al., 2022) affect solubility of the obtained hydrolysate, peptides often show better bulk solubility than plant protein isolates obtained through e.g. heat and/or acid precipitation (Vogelsang-O’Dwyer et al., 2022; Yalcion & Celik, 2007). The solubility of DCH generally displayed a sharp decrease from pH 10 to 12, likely due to alkaline-induced denaturation, which promotes protein aggregates and reduces solubility under the strong alkaline condition. Moreover, DCH displayed high solubility from pH 2 to pH 10, and similarly to DC, a slight decrease in solubility around pH 7. This indicates that a substantial proportion of the protein in DC, and peptides in DCH to an even higher extent, have near-neutral isoelectric points. Overall, pH 8 was shown to be an optimal pH for the CGPs, except for AHP, which was completely insoluble at pH 8. This can be due to aggregation of proteins under the combination of two harsh precipitation methods. The quantitative analysis of pH-dependent protein solubility in the GPCs was also reflected qualitatively by SDS-PAGE analysis (Fig. S1).

**Figure 1.**
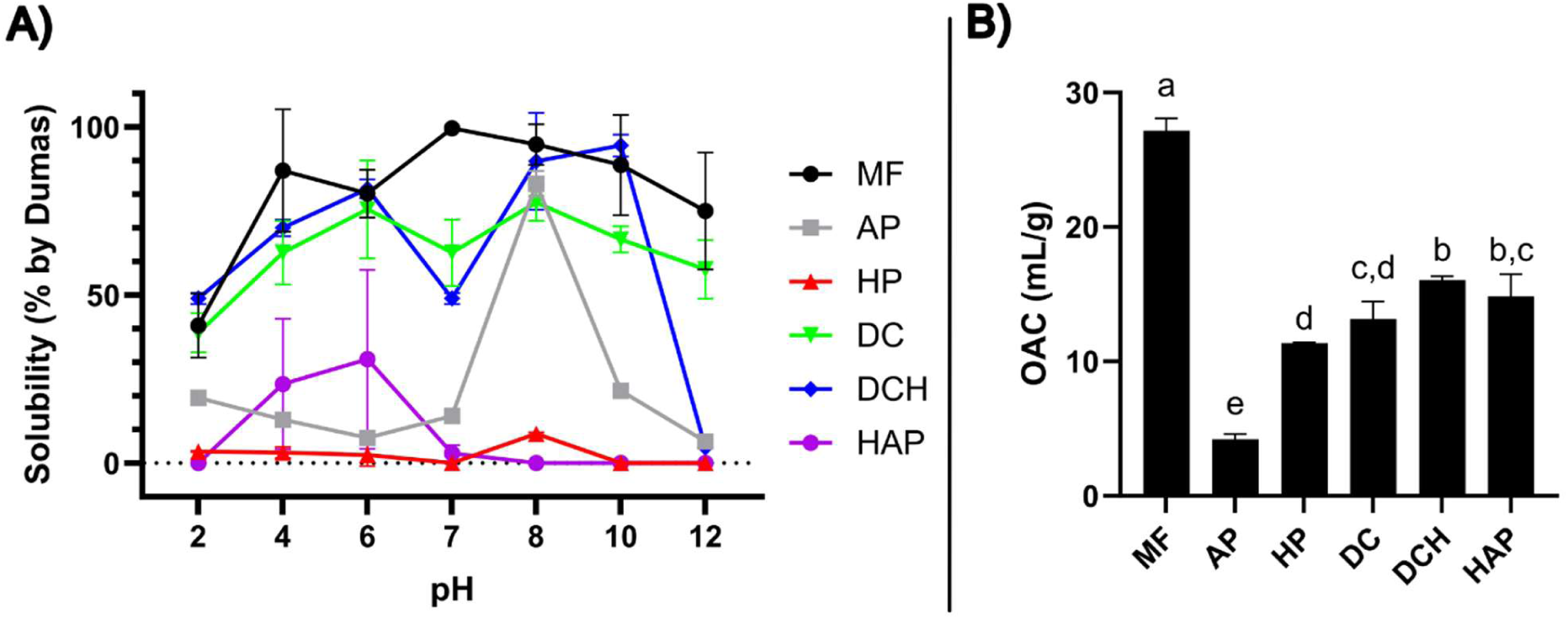
pH-dependent solubility and oil absorption capacity of clover grass protein prototype (CGP) powders. A) Solubility (by Dumas) of prototypes from pH 2 to pH 12. Values are plotted as mean ± SD (n=3). B) OAC of freeze-dried CGP powders. Different letters indicate statistically significant differences between sample means (p<0.05). Values are mean ± SD (n=3). Sample codes: MF: First stage membrane filtration permeate. AP: Acid precipitate. HP: Heat precipitate. DC: Ultra- and diafiltration concentrate. DCH: DC hydrolysate. AHP: Acid and heat precipitate.

#### 3.3.2. Oil Absorption Capacity (OAC)

Oil absorption represents a key functional attribute characteristic of the meat and confectionery industries (Wang et al., 2020). This property relies on the physical entrapment of fat, which is influenced by the hydrophobic surface of protein, protein mass density, and amino acid composition (Ma, 2004). The highest OAC value were obtained for MF (27.16 mL oil/g), whereas AP displayed the lowest (4.21 mL oil/g) OAC (Fig. 1B). HP, DC, HAP, and DCH displayed OAC at comparable levels (11.35-16.02 mL oil/g). Due to the low CP content in MF, and thus high amount of non-protein content, it is not possible to explicitly evaluate if the observed OAC relates to the properties of the protein or other constituents in the prototype. Enzymatic hydrolysis (DCH) demonstrated a significantly (p< 0.05) enhanced ability for absorption compared to the hydrolysis substrate (DC). This improvement is attributed to the cleavage of the peptide bonds in the DCH protein prototype structure, which exposes hydrophobic amino acid residues (such as leucine, isoleucine, and valine). These residues interact favorably with oil molecules, thereby increasing the oil absorption capacity. Similarly, heat-induced denaturation (HP and AHP) appeared to increase surface hydrophobicity of CGP to promote oil absorption.

Interestingly, the combination of acid and heat treatment resulted in higher OAC ability than acid and heat treatment alone, suggesting that the combined heat and acid precipitation may facilitate a higher degree of protein denaturation due to the harshness of the combined methods. The AHP prototype exhibited lower solubility across a wide pH range compared to the other CGPs, indicating that the AHP treatment altered the protein structure in such a way that the hydrophobic chains of proteins were exposed on the surface, allowing them to readily bond with and entrap oil molecules within their structure.

To dive further into the functionality, the results from solubility and OAC analysis were used to delimit the number of prototypes for further analysis, focusing on the prototypes displaying superior solubility and OAC properties (MF, DC, and DCH). This approach allowed us to maintain the integrity of our experimental design while still exploring a diverse range of prototypes. Moreover, two Can protein-based ingredients (pea protein isolate (PPI) and sodium caseinate (Na-Cas)) were included as benchmark references to evaluate the performance of the CGP prototypes.

#### 3.3.3. Properties of selected GCP prototypes at the water-oil interface

Interfacial tension (IFT) was measured using the pendant drop method with 0.1% (w/w) protein concentration as a function of time (Fig. 2A). No reduction in the IFT was observed for the water droplet, confirming the calibrated condition of the tensiometer instrument and the absence of any surface-active compounds in the water. The IFT value remained constant at 26 (mN/m), consistent with previous studies by Badfar *et al*., (2024), Yesiltas *et al*., (2021), and Gultekin Subasi et al., (2022).

**Figure 2.**
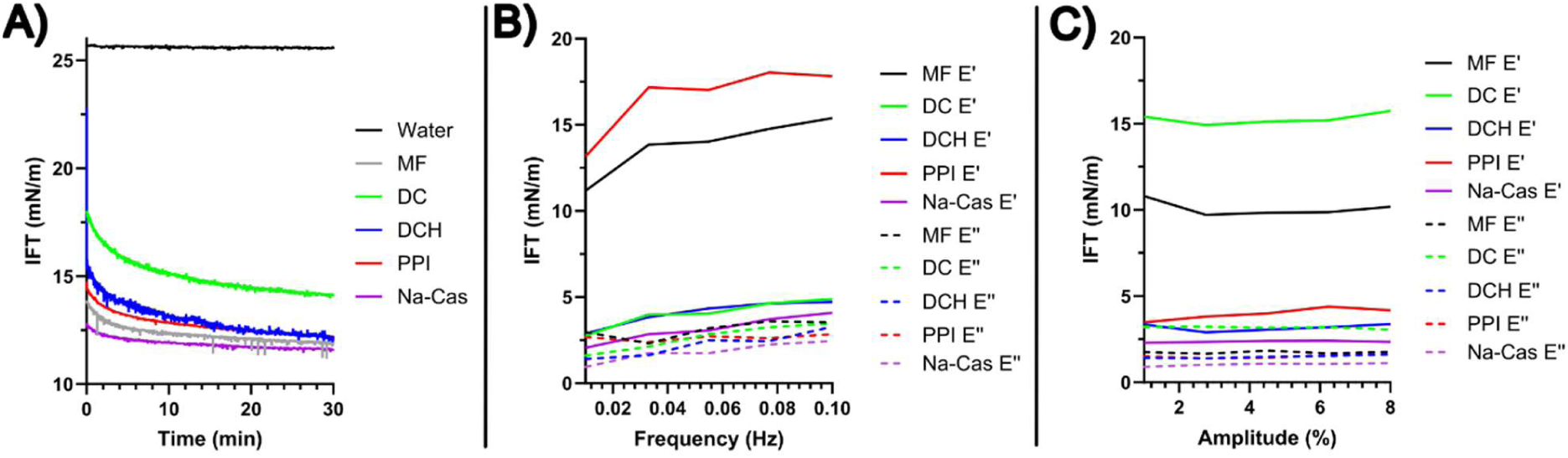
Interfacial tension reduction and dilatational rheology of selected clover grass protein (CGP) prototypes in comparison with a commercial pea protein isolate (PPI) and sodium caseinate (Na-Cas). A) Interfacial tension by pendant drop (water in MCT oil) using 0.1% (w/v) protein solutions over 30 min. B) Elastic (E’) and viscous (E’’) moduli from the frequency sweep test from 0.01 Hz to 0.1 Hz. C) Elastic (E’) and viscous (E’’) moduli from the amplitude sweep test from 1% to 8% amplitude. Sample codes: MF: First stage membrane filtration permeate. DC: Ultra- and diafiltration concentrate. DCH: DC hydrolysate.

The lowest initial (13 mN/m) and final (12 mN/m) IFT values were achieved by Na-Cas, though all three CGPs displayed ability to reduce IFT over 30 min. DC exhibited the lowest capability in reducing IFT both at 0 min and 30 min, whereas the hydrolysate produced here from (DCH) demonstrated a substantially improved ability to reduce IFT. DCH showed a sharp initial decrease, indicating rapid protein adsorption at the interface (Amine et al., 2014). This reduction was followed by a gradual decline, reflecting conformational rearrangement at the water/oil interface (Amine et al., 2014). The lower reduction (higher value) of IFT for DC prototype could be attributed to its lower solubility compared to DCH and MF at pH 7, indicating less soluble protein available for absorption at the water-oil interface. Furthermore, the MF prototype exhibited superior IFT reduction capability among the other CGPs and PPI, suggesting that fewer processing steps altered the clover grass protein conformation in a way that affected their amphiphilic nature. Consequently, MF with one or two fewer processing steps than DC and DCH, demonstrated a better ability to reduce the IFT value initially and after 30 min. Nevertheless, the low CP content in MF makes it difficult to say whether improved IFT reduction capacity relates to protein state or non-protein components in the prototype. The ability of plant protein sources to reduce IFT has been studied previously. Tan et al., (2022) reported that the RuBisCO protein extracted from duckweed (*Lemna minor*) exhibited an IFT of approximately 15 mN/m, which was lower than the IFT of soy isolate protein (≈16 mN/m), a plant protein, but higher than whey protein (≈14 mN/m), an animal protein source. The equilibrium IFT values (30 min) obtained from all three CGPs and reference proteins were comparable or lower to these values, indicating a comparable or better ability to stabilize the oil/water interface.

Analysis by dilatational rheology reveals information on the resistance to surface deformation of an interface and can help determine the mechanical properties of the interfacial layer (McClements et al., 2022). Hence, this study aimed to elucidate the dilatational rheological properties of CGPs and compare their molecular behavior at the O/W interface with controls (Na-Cas and PPI) at a concentration of 0.1% (w/w). The exposed amino acids interact intermolecularly with adjacent protein chains at the interface, forming a viscoelastic film at the O/W interface. The characteristics of this viscoelastic film reveal information about the ability of the interfacial protein layer to reduce IFT (Kontogiorgos & Prakash, 2023). The frequency sweep test was performed within a range of 0.01-0.1 Hz with a constant amplitude of 4% to assess the response of each CGP and controls to increasing frequency over time (Fig. 2B). According to the results, E’ (elastic modulus) values dominated the E’’ (viscous modulus) in all three CGPs and controls, reflecting the viscoelastic nature of the interfaces (Badfar et al., 2022). The patterns of E’ and E’’ exhibited slight dependency on the increase of the frequency. Minimal frequency dependence signifies that the interfacial films are well-structured, as alterations in the surface area (ΔA/Ao) do not affect relaxation processes over either short (high frequency, rapid relaxation times) or long (low frequency, slow relaxation times) durations. Comparing the samples, DC and MF-treated CGPs displayed the highest E′ values, indicating that they formed the most elastic and rigid O/W interfacial layers. Their strong elastic response to frequency changes was even greater than that observed for DCH, Na-Cas, and PPI. Since higher E′ values correspond to a more solid-like interface, whereas lower E′ values reflect a more flexible or stretchable interfacial film (Tamm & Drusch, 2017), these results highlight the distinct rigidity of the DC and MF-treated interfaces. In contrast, sample Na-Cas showed consistently low E′ values that were unaffected by frequency, which aligns with previous studies (Badfar et al., 2025; Gultekin Subasi et al., 2022b), and suggests that Na-Cas forms a more stretchable, less elastic interface. All samples exhibited relatively low viscous moduli (E″), with sample DC showing the highest energy dissipation. Under linear frequency conditions, the low E″ values indicate limited viscous losses during dilatational cycling. Nevertheless, increases in E″ are generally associated with greater interfacial mobility and the presence of dissipative mechanisms such as adsorption–desorption, diffusional relaxation, or conformational rearrangements, along with possible hydrodynamic coupling to the subphase (Risse et al., 2025).

The amplitude sweep test was carried out, while the frequency was maintained at a constant value of 0.1 Hz (Fig. 2C). All samples exhibited a linear viscoelastic range (LVR) within the deformation range of 0.1-0.3 mm. With increasing amplitude (%) and subsequent deformation, E’ value began to fluctuate slightly in both CGPs and controls. These fluctuations in the E’ tended to decrease for DC, DCH, and MF prototypes. However, this trend was not observed in the controls, where E’ remained nearly constant for Na-Cas and increased for PPI. These behaviors are consistent with previous interfacial dilatational studies, where stronger and less-stretchable films showed stable or rising E′ with amplitude, while more reconfigurable films soften as deformation grew (Badfar et al., 2025; García-Moreno et al., 2021). The highest elastic element was recorded for DC, demonstrating a stronger and more elastic-like structure at the O/W interface compared to the other CGPs and controls. A stable and constant response to the increased deformation rate suggests a more stretchable (solid-like) but weaker interface structure (Yang et al., 2020). A similar viscoelastic response for Na-Cas was reported previously (Subas et al., 2022; Badfar et al., 2024). The diminished interactions at oil interfaces can be readily explained by the fact that hydrophobic amino acids exhibit a higher affinity for oil, consequently reducing interactions between amino acids (Kontogiorgos & Prakash, 2023).

Two aspects regarding IFT and dilatational rheology experiments are necessary to consider. Firstly, it is important to note that the protein concentrations used in these tests are lower than those typically found in food applications (i.e. 0.1% in this study). Consequently, phenomena such as protein aggregation may not be manifested under these conditions. Secondly, dilatational rheology is performed on a single droplet, which does not accurately represent the numerous smaller droplets present in real emulsions. This discrepancy means that protein interactions on the droplet surfaces, which could lead to potential flocculation and stability issues, were not captured by this method (Pérez-Vila et al., 2024). Therefore, it is crucial to further investigate the emulsifying properties of the proteins in actual emulsion systems.

### 3.4. Physical stability of emulsions stabilized by selected CGPs

Various plant sources, including legumes and cereals, can provide surface-active proteins (McClements, 2015). Many investigations have been conducted to examine the potential of these proteins to stabilize emulsions. Following a two-stage preparation of oil-in-water emulsion (5%, v/v), the ζ-potential after Day 1 and Day 8 of storage was recorded (Table 3). On Day 1, the ζ-potential of PPI, serving as a plant control, was recorded at a significantly lower (p<0.05) absolute value than that of CGPs and Na-Cas. Sufficient repulsion, indicated by an absolute ζ-potential value greater than 30 mV, is necessary to prevent droplet merging (McClements, 2015). This suggests that the repulsion between PPI emulsion droplets was insufficient to prevent droplet coalescence. A ζ-potential absolute value of 24.6 mV for the emulsion stabilized by pea protein isolated (PPI) has been previously reported by (Sha & Xiong 2022), and aligns with our finding of 27.82 mV. CGPs exhibited a ζ-potential value comparable to Na-Cas (p>0.05). Moreover, the absolute ζ-potential values for CGPs and Na-Cas increased from Day 1 to Day 8. This change was particularly pronounced in the MF and DC prototypes. A possible explanation for this observation is potential oxidation processes in the oil content, which releases chemical compounds, causing a rearrangement of the interfacial composition, subsequently leading to physical destabilization of the emulsion system (Vall-llosera et al., 2021). Na-Cas, as a positive control, showed a ζ-potential value of -43.20 mV on Day 1 of storage, consistent with findings (Badfar et al., (2025), Yesiltas et al., (2021)).

**Table 3.** Droplet sizes (D(4,3) and D(3,2)) and ζ-potential of 5% emulsion stabilized by 0.2% clover grass protein prototypes (CGPs), pea protein isolate (PPI), or sodium caseinate (Na-Cas).

| Sample | Droplet sizes ($\mu\text{m}$) | | | | | | ζ-potential (mV) | |
| --- | --- | --- | --- | --- | --- | --- | --- | --- |
|  | Day 1 |  | Day 3 |  | Day 8 |  | Day 1 | Day 8 |
|  | D(4,3) | D(3,2) | D(4,3) | D(3,2) | D(4,3) | D(3,2) |  |  |
| MF | 24.28±1.63<sup>a</sup> | 8.69±0.67<sup>a</sup> | 13.89±2.08<sup>b</sup> | 5.30±0.23<sup>a</sup> | 28.03±0.15<sup>a</sup> | 2.00±0.01<sup>a</sup> | -44.05±0.45<sup>a</sup> | -57.30±1.92<sup>b</sup> |
| DC | 7.94±0.66<sup>b</sup> | 1.99±0.01<sup>c</sup> | 5.90±0.35<sup>c</sup> | 1.91±0.00<sup>c</sup> | 3.56±0.01<sup>d</sup> | 1.43±0.04<sup>bc</sup> | -46.20±1.66<sup>a</sup> | -65.60±2.00<sup>a</sup> |
| DCH | 21.30±1.90<sup>a</sup> | 2.96±0.14<sup>b</sup> | 24.47±2.28<sup>a</sup> | 2.88±0.10<sup>b</sup> | 18.26±0.68<sup>b</sup> | 1.28±0.06<sup>cd</sup> | -43.45±3.09<sup>a</sup> | -48.60±2.45<sup>c</sup> |
| PPI | 3.50±2.30<sup>c</sup> | 0.57±0.03<sup>d</sup> | 3.35±0.23<sup>cd</sup> | 0.64±0.03<sup>d</sup> | 4.29±0.10<sup>c</sup> | 1.51±0.01<sup>b</sup> | -27.82±1.03<sup>b</sup> | -27.02±1.53<sup>d</sup> |
| Na-Cas | 0.76±0.03<sup>c</sup> | 0.38±0.01<sup>d</sup> | 0.83±0.83<sup>d</sup> | 0.37±0.36<sup>e</sup> | 2.97±0.07<sup>d</sup> | 1.14±0.15<sup>d</sup> | -43.20±1.35<sup>a</sup> | -45.32±3.69<sup>c</sup> |

Different letters within the same column indicate significant differences between means ( $p < 0.05$ ). Values are mean ± SD (n=3).
Sample codes: MF: First stage membrane filtration permeate. DC: Ultra- and diafiltration concentrate. DCH: DC hydrolysate.

Droplet size measurement can provide critical insights into the emulsifying properties of CGP prototypes. In the current study, droplet sizes were determined by volume-weighted mean (D4,3) and surface-weighted mean (D3,2) in 5% O/W emulsions stabilized with 0.4% CGPs on Day 1, 3, and 8 (Table 3). The positive control Na-Cas exhibited the lowest D4,3 and D3,2 on Day 1, 3, and 8 of the storage period. The plant-based control (PPI) appeared to produce smaller droplets than CGP prototypes, but still significantly larger than Na-Cas (p<0.05). Similar findings for Na-Cas have previously been reported by Yesiltas et al. (2021). Additionally, Sha et al. (2022) documented the droplet size of pea protein emulsion (2.01 µm) produced using sunflower oil and 10 mg/ml protein in phosphate buffer. However, their measurement was based on the intensity-weighted Z-average hydrodynamic diameter (DLS, Zetasizer Nano S90, 90°).

Conversely, the MF and DCH prototypes displayed a larger volume of droplets and a lower number of small droplets compared to DC and the controls during the storage period. These observations suggest that, despite having a high ζ-potential, CGPs produced larger oil droplets compared to the controls. This observation indicates that emulsions stabilized with 0.4% CGPs are prone to phase separation or coalescence, potentially compromising their physical stability. Among the CGP prototypes, DC produced the smallest droplets. Furthermore, the droplet size for DC-emulsions reduced over time to a level comparable to that of both PPI and Na-Cas on Day 8. The decrease in the droplet sizes of CGP emulsions may relate to the creaming in the resulting emulsions. Such a phenomenon has been reported previously by Martin *et al*. (2019) and Pérez-Vila *et al*. (2024).

It may appear that with 0.4% CGPs, concentration was insufficient to adequately stabilize the surface of the oil droplets, resulting in larger droplet sizes (McClements et al., 2022). The reduction in droplet size with increasing protein concentration from 0.2 to 1% (w/w) has previously been reported by Pérez-Vila *et al*. (2024) for 10% sunflower emulsions stabilized with ryegrass protein. Under certain conditions, such as at specific pH levels, oil droplets can be covered with a higher amount of protein due to protein aggregation, leading to a thicker protein layer and, consequently, larger droplets.

Immediately after production, the emulsion stabilized with DCH displayed no visible creaming (Fig. 3), suggesting that DCH, as an emulsifier at the oil-water interface, can provide initial stabilization, preventing immediate coalescence or aggregation of droplets. However, by Day 1, all the emulsions stabilized with CGPs exhibited instability and showed phase separation, which is commonly due to the differences between the continuous and dispersed phase density. Based on Stoke’s law, the creaming velocity increases if the droplet radius and oil droplet size are large (Loi et al., 2019; Wilde, 2019). Compared with the ζ-potential results, CGPs were expected to exhibit reduced creaming formation, as they showed sufficient repulsion between the droplets. However, the PPI emulsion displayed smaller droplets (Table 3) and no visible creaming until Day 6 of storage (Fig. 3). The emulsion stabilized with Na-Cas was the most stable during the storage period, displaying the smallest degree of creaming with onset on Day 6. The most pronounced creaming was observed in the MF prototype. As previously mentioned, the content of non-protein constituents (including added buffer salts) may have a large effect on in vitro functionality of this particular prototype. Overall, the results of physical stability tests confirmed that the ability of CGPs at a concentration of 0.4% (w/w) to stabilize the rapeseed oil in water emulsion is not favorable.

**Figure 3:**
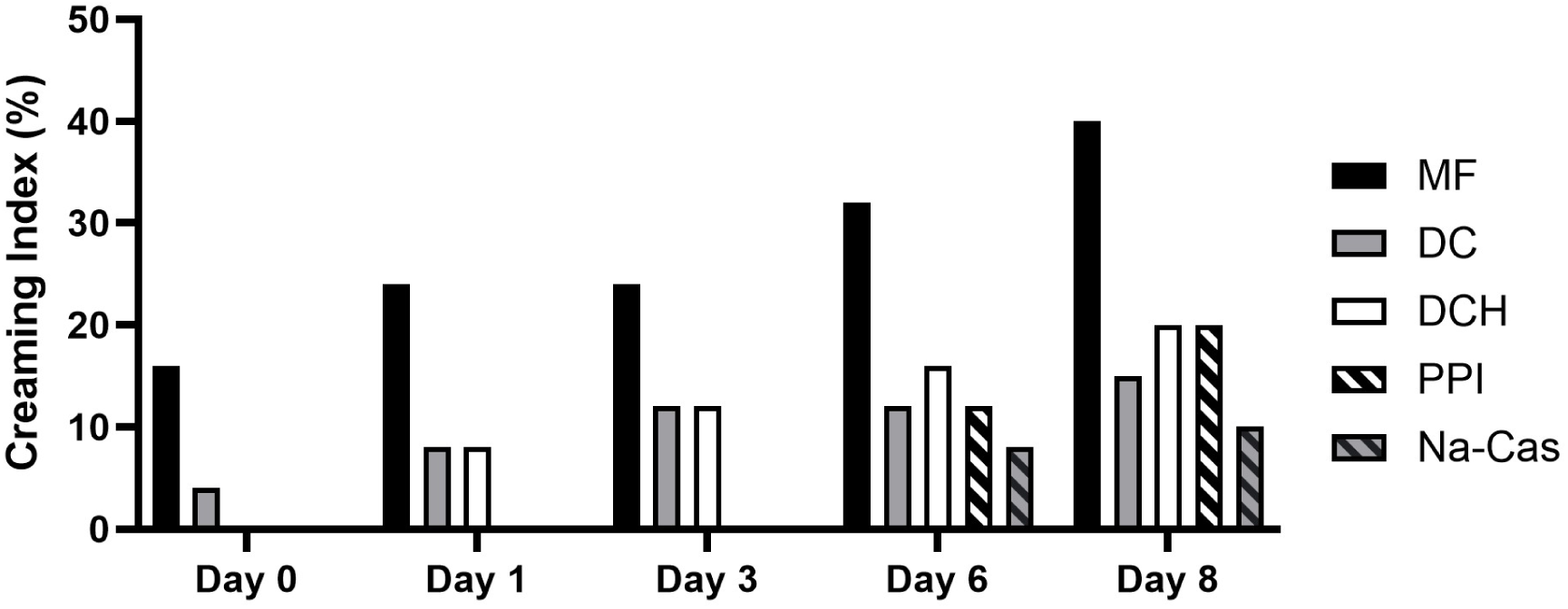
Creaming index for 5% rapeseed oil emulsions stabilized by 0.4% (w/v) clover grass protein prototypes (CGPs), pea protein isolate (PPI), and sodium caseinate (Na-Cas) over eight days of storage. Sample codes: MF: First stage membrane filtration permeate. DC: Ultra- and diafiltration concentrate. DCH: DC hydrolysate.

### 3.5. Analysis of differential protein composition by label-free quantification

A total of 2018 proteins were identified across all samples, whereof 1740 were reproducibly quantified in at least one stream/prototype. Analyzing the intersection of reproducible IDs across streams/prototypes (Fig. 4A), the most frequent observation was that a protein (388 or 22%) was identified in all streams. Moreover, a quite large proportion (174 or 10%) were only identified in the initial green juice Feed stream, while proteins not observed in DCH constituted 115 proteins or 6.6% of all reproducibly identified proteins. Feed-specific proteins can be ascribed to the selective retention of certain proteins in the first-stage membrane operation, as previously observed (Gregersen Echers et al., 2026; Mattsson et al., 2025). In contrast, selective depletion in the hydrolysate cannot be explained by membrane selectivity but may rather be ascribed to either certain proteins being resistant to *in vitro* tryptic hydrolysis or by substrate-level selectivity according to classical Linderstrøm-Lang theory, as hydrolysis was not allowed to progress to completion (Christensen et al., 2025). This theory states that a protease is more inclined to fully hydrolyze one protein before initiating hydrolysis of the next (Linderstrom-Lang K., 1953). Another potential explanation is that some proteins may produce hydrophobic, thermo-labile or aggregation-prone peptide products, which would be precipitated in the heat-induced hydrolysis termination. Surprisingly, the MF prototype, which was the basis for developing the remaining downstream prototypes, had a substantially lower number of IDs compared to both Feed and remaining prototypes, when disregarding DCH (Fig. 4A). A full overview of overlapping protein IDs between the Feed and the prototypes can be found in the supplementary information (Fig. S2).

**Figure 4:**
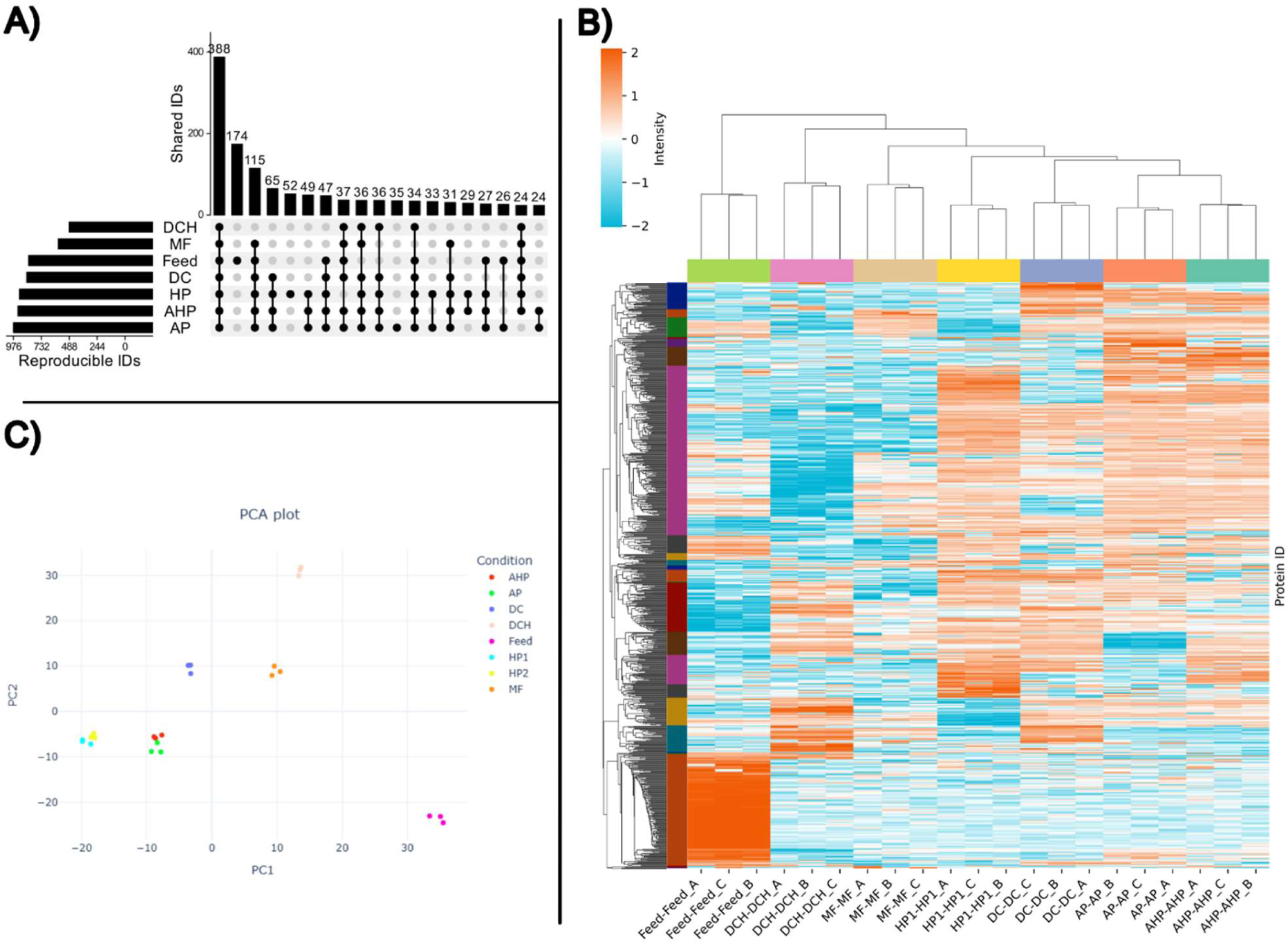
Qualitative and MaxLFQ-based quantitative analysis of LC-MS/MS data from the green juice Feed stream and all clover grass proteins (CGPs). A) UpSet plot showing the number and overlap of reproducible (quantified in at least two of the replicates) protein IDs across the feed and GCP prototypes. Only intersects with more than 20 protein IDs are shown. For a full visualization of all intersections, please refer to the supplementary information (Fig. S2). B) Heatmap representation of the 835 differential proteins from ANOVA-based analysis across the Feed stream and all CGP prototypes. Data is depicted as z-score standardized MaxLFQ intensities by row (protein group) and clustered using a Euclidian distance of 5. C) Principal component analysis (PCA) showing replicate-wise sample similarity and clustering according to overall qualitative and quantitative variability. Individual sample replicates are depicted based on the two first principal components, explaining 14.5% and 11.5% of the total variability, respectively.

That the initial Feed stream stood out the most from the prototypes is also evident when taking into account LFQ-based quantification in an ANOVA-based differential analysis. In total, 25 protein-level clusters were observed using a Euclidean distance of 5. (Fig. 4B). While the large number of clusters makes interpretation challenging, increasing cluster distance only produced four clusters, which were considered inadequate at visually describing the variability in the data set (data not shown). One of four major clusters (Fig. 4B, red, bottom) represents 159 proteins that appear highly enriched in the feed stream. This is in good agreement with the qualitative analysis (Fig. 4A). The largest cluster represents 242 proteins (Fig. 4B, purple, top), that generally appear highly depleted in DCH. This cluster therefore not only represents the 155 proteins not reproducibly identified in DCH (Fig. 4A) but also additional proteins that were of substantially lower abundance, further substantiating the potential selectivity imposed by *in vitro* hydrolysis. Overall, ANOVA-based differential analysis revealed that there are indeed distinct quantitative differences between prototypes and based on sample-level clustering, AP and AHP appeared the most similar while DCH differentiates most from the rest of the prototypes and Feed is the most differentiation sample among all.

This differentiation is also reflected by the PCA analysis performed (Fig. 4C). Here, the feed stream differs from the prototypes on both PC1 and PC2, while DCH is more similar to the rest on PC1 but differs substantially on PC2. Similarly, AP and AHP co-localize in the biplot. Using the MF prototype as a starting point for producing the remaining prototypes, it appears that DC is only affected along PC1. AP and AHP are further affected along PC1 but also along PC2, while HP is affected even more along PC1 but along PC2 to the same extent as AP and AHP. As such, it could appear that while AHP is subjected to both acid and heat, it is acid treatment that is dominant to the protein composition in the prototype. This is an interesting observation, as for functionality aspects, it appears that AHP has properties closer to HP than AP when considering solubility and OAC. As such, the application of acid appears to largely affect the protein composition while the application of heat has a more pronounced effect functionality. As such, the observations indicate that protein state (i.e. heat denatured) appears to be more important for determining functionality than protein-level compositional differences. Nevertheless, the magnitude of the observed differences and the effect on the bulk protein composition is not evident from LFQ-based analysis. Consequently, we applied a secondary and complementary quantification using relative intensity-based absolute quantification (riBAQ) to identify process-induced effects on the most abundant proteins across the prototypes.

### 3.6. Identification of abundant proteins by iBAQ quantification

Across the Feed stream and all analyzed CGP prototypes, 63 proteins were determined as highly abundant (mean riBAQ > 0.5%) in at least one sample (Fig. 5A). Among the most abundant proteins, RuBisCO (rbc) subunits were not surprisingly the most dominant. Of the 63 abundant proteins, 54 were found as highly abundant in a previous study using the same abundance threshold (Gregersen Echers et al., 2026). While this represents a generally high degree of overlap between studies, the observed differences may be ascribed to variables such as time of harvest, growth stage and biomass composition. While biomass was obtained from the same field as in the previous study, differences in composition, both in terms of biomass composition and protein content within the individual plants, are bound to occur across a season. The nine additional abundant proteins covered several stress response proteins (abr17, PR-10, and PR-4), which may be related to soil/plant health at time of harvest. While all nine are below 0.6% in the Feed stream, and eight are maintained at levels below 1% in any of the prototypes, PR-10 becomes significantly enriched (p < 0.05) in all prototypes but HP. In fact, in AHP, it is 10-fold enriched to 2% from the initial 0.2% in the Feed stream. This is relevant as sensitization to PR-10 pollen proteins, as well as other abundant and related protein like Bet v 1, from e.g. grasses or birch, may lead to primary sensitization or cross-reactive allergenic response in certain foods. nsLTPs, the well-known food allergens with sensitization capabilities, were in our previous study reported to become enriched after the first stage filtration but subsequently almost fully depleted after ultra- and diafiltration (Gregersen Echers et al., 2026). Here, a 2.5-fold and significant (p < 0.0001) enrichment (0.2% to 0.5%) across the initial filtration stage was also observed, but in all prototypes except for HP (0.2%), the nsLTP content was minimal (< 0.03%) and significantly lower (p < 0.0001) as well as 10-fold lower than in DC from our previous study (0.2%) (Gregersen Echers et al., 2026).

**Figure 5:**
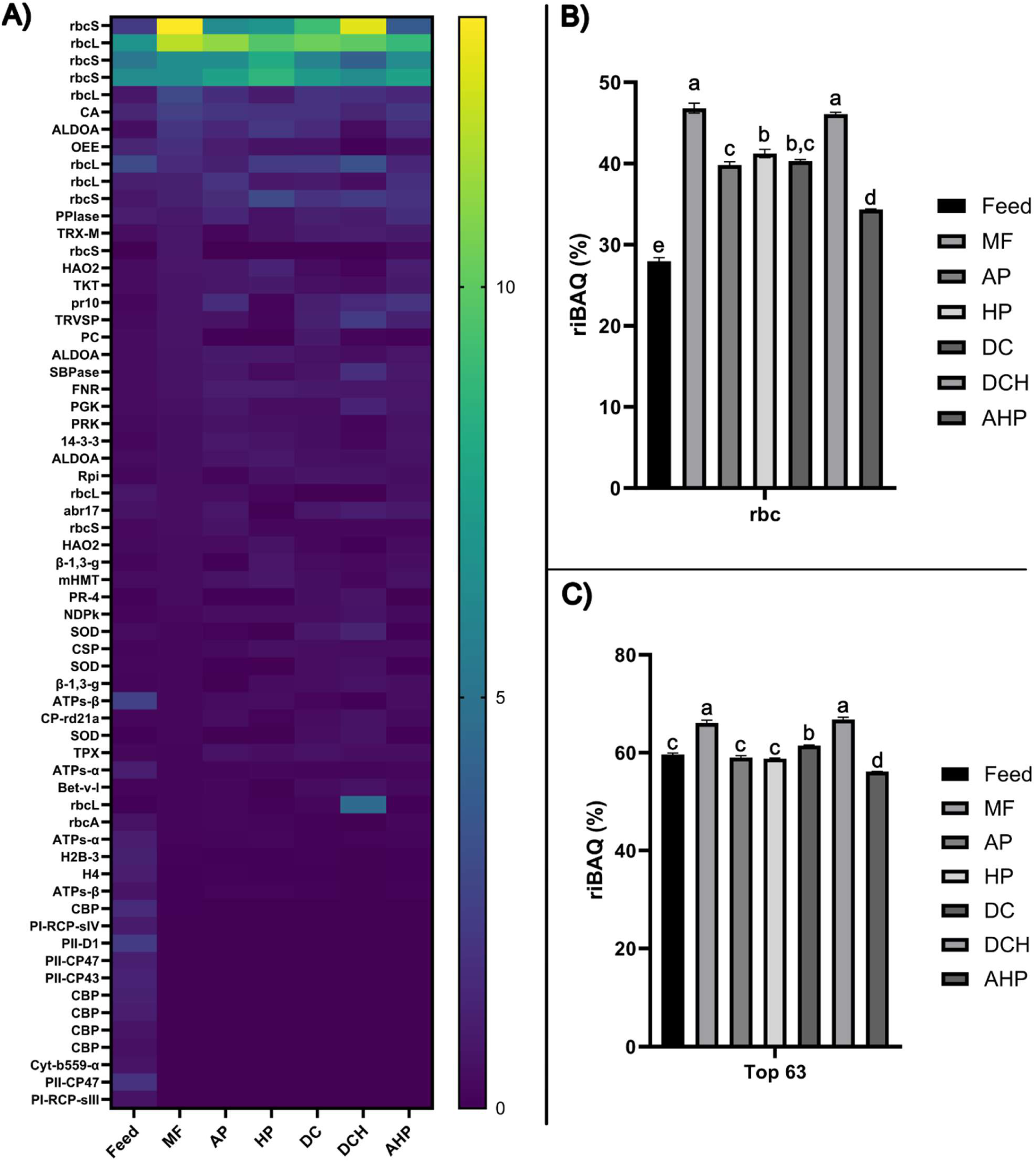
Differential protein abundance across the Feed stream and CGP prototypes by relative iBAQ quantification. A) Heatmap of mean riBAQ for proteins found above a 0.5% riBAQ threshold (top63) in any of the samples. Mean protein abundance (by % riBAQ) is color coded from low (blue) to high (yellow) abundance. Proteins are given using short names as defined in Table S2. B) Total RuBisCO (rbc) abundance (sum of all rbc subunits and isoforms) across the initial Feed stream and all CGP prototypes. Abundance is indicated as ΣriBAQ (mean +/- SD) based on triplicate analysis for each sample. C) Cumulative abundance of all abundant (top63) across the initial feed stream and all CGP prototypes. Abundance is indicated as ΣriBAQ (mean +/- SD) based on triplicate analysis for each sample. In histogram plots (B and C), letters above each column indicates statistical similarity (p < 0.05) in a group-wise manner based on one-way ANOVA with Tukey.

The initial feed stream has a rbc content of 28%, which is substantially and significantly (p < 0.0001) enriched in the MF permeate stream to 47% (Fig. 5B). This illustrates the ability of the first crossflow filtration step to increase the relative proportion of rbc in the permeate. In a separate study on green juice from isolated and lab-grown *L. perenne*, the rbc content was found to be 35-39% (Danner Aakjaer Pedersen et al., 2025), indicating that growth conditions in a climate chamber may be more suitable for obtaining high rbc leaves or that the other species in the mixed field may contribute with a lower rbc content to the bulk biomass. While rbc content was here significantly (p < 0.0001) depleted in AP, HP, DC, and AHP, high levels (34-46%) are retained in the different downstream prototypes (Fig. 5B). The initial rcb content in green juice is comparable to our previous study on biomass from the same field finding 26% rbc, the content is the permeate stream after initial crossflow filtration is substantially higher than the previous study finding 19%. This highly substantial and more than 2.4-fold increase in rbc content may be ascribed to a better performing process for this particular batch than in earlier work. Moreover, this is also reflected by the rbc content in DC (40%), where the previous study found 17%. Chlorophyll a-b binding proteins (CBP) and Photosystem I & II (PI/II) proteins were principally fully depleted following the initial membrane filtration (Fig. S3). This is in agreement with our previous study, where we found that the initial crossflow membrane process is capable of clearing green juice by selective retention of pigment-binding and membrane-associated proteins (Gregersen Echers et al., 2026).

While some specific proteins like PR-4, plastocyanin, and superoxide dismutase appeared to be more selective depleted with application of heat and/or acid to precipitate proteins (Fig. 5A), the overall quantitative protein composition in the prototypes appeared quite similar. Considering how much of the total protein is constituted by the 63 most abundant proteins combined, the difference between prototypes is not tremendous. Top 63 represents 59.6% of the total protein in the initial stream (Feed) and is enriched to 66.1% in the first stage membrane filtration (MF) permeate (Fig. 5C). This is despite the selective retention of abundant pigment-binding and membrane-associated proteins on the membrane. This is, in turn, compensated for by a significant increase in rbc content (Fig. 5B). The level of top 63 proteins is retained in DCH (66.8%) but depleted slightly in the remaining prototypes (56.1%-61.5%).

While AP and AHP were more similar using LFQ-based quantification (Fig. 4), AHP was closer to HP in terms of functionality (Fig. 1, Fig. 2). Considering the overall abundance distribution of top 63 proteins (Fig. 5A), AHP appears more similar to AP than to HP, corroborating the LFQ-based analysis. However, when considering the bulk abundance of the primary constituent rbc (Fig. 5B) as well as the cumulative abundance of top 63 (Fig. 5C), AP and HP resemble each other more than they resemble AHP. As such, it is not seemingly possible to couple the protein-level composition to the differences observed in the bulk properties of the prototypes. Rather, the observed differences are more likely to be ascribed to the physical state of the protein. For instance, acid, and particularly heat, treatment dramatically reduced protein solubility across a broad pH range compared to the native protein in DC. A lower aggregation state in more native prototypes seems to be associated with an initially better solubility in the aqueous phase, as previously reported for spinach rbc but also seen in e.g. dairy protein. The aggregation state of plant proteins has previously been demonstrated to affect interfacial adsorption behavior and emulsifying properties. However, increased aggregation state may be beneficial for faster adsorption and more efficient interface stabilization in rbc.

## Conclusion

As the hunt for sustainable, plant-based protein food ingredients intensifies, green biomasses, such as grasses and clovers, are often highlighted as sources of great potential. This is particularly based on the high content of RuBisCO, which has a good nutritional profile and promising functional properties. However, the process in which such a protein ingredient can be obtained has not fully been established. A major bottleneck is the fact that methods for protein isolation have a substantial effect on the protein-level composition, yields, and bulk properties of a protein product. As such, the current study focused on both techno-functional and physiochemical properties as well as protein-level composition of clover grass protein (CGP) obtained using different processing techniques for a clarified green juice. The methods covered precipitation by acid (AP), heat (HP), or combined (AHP) as well as a gentler method using membrane-base ultra- and diafiltration (DC) and subsequent tryptic hydrolysis (DCH). Results revealed that different processing methods significantly affect CGP characteristics. HP exhibited the highest protein content (69%), while the initial clarified juice from membrane filtration (MF) had a very low protein content (8.7%). The remaining CGP prototypes displayed similar protein content (51-57%). All CGP prototypes showed pH-dependent solubility, but the protype obtained from pure membrane-based methods (DC) and its hydrolysate (DCH) displayed much higher solubility across a wide pH range, in contrast to the prototypes obtained from heat precipitation (HP) and combined acid and heat precipitation (AHP) which displayed low solubility. Combined heat and acid treatment resulted in the highest protein purity and the whitest prototype, however at the cost of impaired functionality. CGPs showed large variation in oil absorption capacity with the highest OAC observed in the initial prototype (MF). While MF, DC, and DCH were all able to reduce O/W interfacial tension in follow-up experiments, these prototypes displayed different viscoelastic responses to amplitude and frequency sweep changes, indicating that different processing methods influenced viscoelastic responses. Among all CGPs, DCH showed a lower elasticity in both amplitude and frequency tests. The application of MF, DC, and DCH as an emulsifier was assessed in the oil-in-water emulsion, and despite possessing a higher absolute value of ζ-potential, CGPs-stabilized emulsions showed larger droplets compared to pea protein and sodium caseinate over an 8-day storage period. The creaming index confirmed that at 0.4% concentration (w/w), the ability of CGPs to stabilize a 5% rapeseed oil in water emulsion is not favorable. Using bottom-up proteomics, the protein composition of the prototypes and the green juice used as feed to produce the initial MF prototype revealed substantial and quantitative differences. While all prototypes revealed a high content of RuBisCO (37-47% of the total protein), which was enriched compared to the green juice (28%), distinct process-induced differences in the abundance of e.g. potential allergen proteins were observed. The analysis revealed that the observed differences in functionality could not be described by differences in the quantitative, protein-level composition but were more affected by the state of the protein, as prototypes with similar protein-level composition displayed substantial differences in functional properties. Ultimately, this study highlights the importance of downstream processing on not only protein-level composition but particularly techno-functional properties of plant proteins for use as food ingredients.

## Author contributions

NB: Methodology, Validation, Formal analysis, Investigation, Writing – original draft preparation, Writing – review and editing, Visualization. SGE: Conceptualization, Methodology, Validation, Formal analysis, Investigation, Writing – original draft preparation, Writing – review and editing, Visualization, Supervision. CJ: Resources, Writing – review and editing, Supervision. BY: Resources, Writing – review and editing, Supervision. AKJ: Methodology, Formal analysis, Investigation, Writing – review and editing. TM: Methodology, Formal analysis, Investigation, Writing – review and editing. PSL: Conceptualization, Funding acquisition, Writing – Review & Editing. AISV: Methodology, Formal analysis, Writing – review and editing. AM: Methodology, Formal analysis, Writing – review and editing. KLB: Conceptualization, Funding acquisition, Writing – Review & Editing. ML: Conceptualization, Methodology, Funding acquisition, Writing – Review & Editing, Supervision.

## Acknowledgments

The authors would like to thank Inge Holmberg and Lis Berner from the Research Group for Bioactives - Analysis and Application (Technical University of Denmark) for their kind assistance. Harvesting and wet fractionation of clover grass was carried out in close collaboration <u>with the</u> Site Manager, Master Machinist Morten Olsen, (BiomassProtein ApS (Denmark)), who also assisted in membrane filtration of green juice to produce the prototypes for this research. The authors would like to extend their gratitude for this assistance.

## Data Availability

The mass spectrometry proteomics data have been deposited into the ProteomeXchange Consortium via the PRIDE partner repository with the dataset identifier PXD070032 and DOI 10.6019/PXD070032. All other data will be made available upon request.

