## Supplementary information for "Techno-functional properties of clover grass protein and the effect of different process technologies"

For

### Content of supplementary material

#### Supplementary tables

**Table S1:** MaxQuant output data and additional downstream processing hereof.

**Table S2:** Overview of the most abundant (riBAQ > 0.5%) proteins in any of the analyzed samples.

#### Supplementary figures

**Figure S1:** Composition of **s**oluble protein at different pH (from solubility test) by SDS-PAGE.

**Figure S2:** Full UpSet Plot from Fig. 4A.

**Figure S3:** Differential abundance (by riBAQ) for nsLTPs, CBPs, and PI/II proteins across all samples.

**Table S1:** MaxQuant output data and additional downstream processing hereof. The “proteinGroups” folder has been modified to include a range of additional data used in the manuscript covering calculation of relative iBAQ (riBAQ), replicate means, abundance and reproducibility filters. For a full data file with no modification, see the referenced project in the PRIDE data repository. Table can be found in appended .xlsx file (Supplementary Tables) as “Table S1”.

**Table S2:** Overview of the most abundant (riBAQ > 0.5%) proteins in any of the analyzed samples. The table includes the Uniprot AC#, protein name, a short name (as used in Figure 5), various MaxQuant Data (number of proteins in group, peptide IDs, unique peptides, sequence coverage, molecular weight and Andromeda score), mean riBAQ abundance across all sample types, The table contains both individual proteins (Maxquant proteinGroups by lead protein ID) and the defined “families/types/groups” from this work. Table can be found in appended .xlsx file (Supplementary Tables) as “Table S2”.

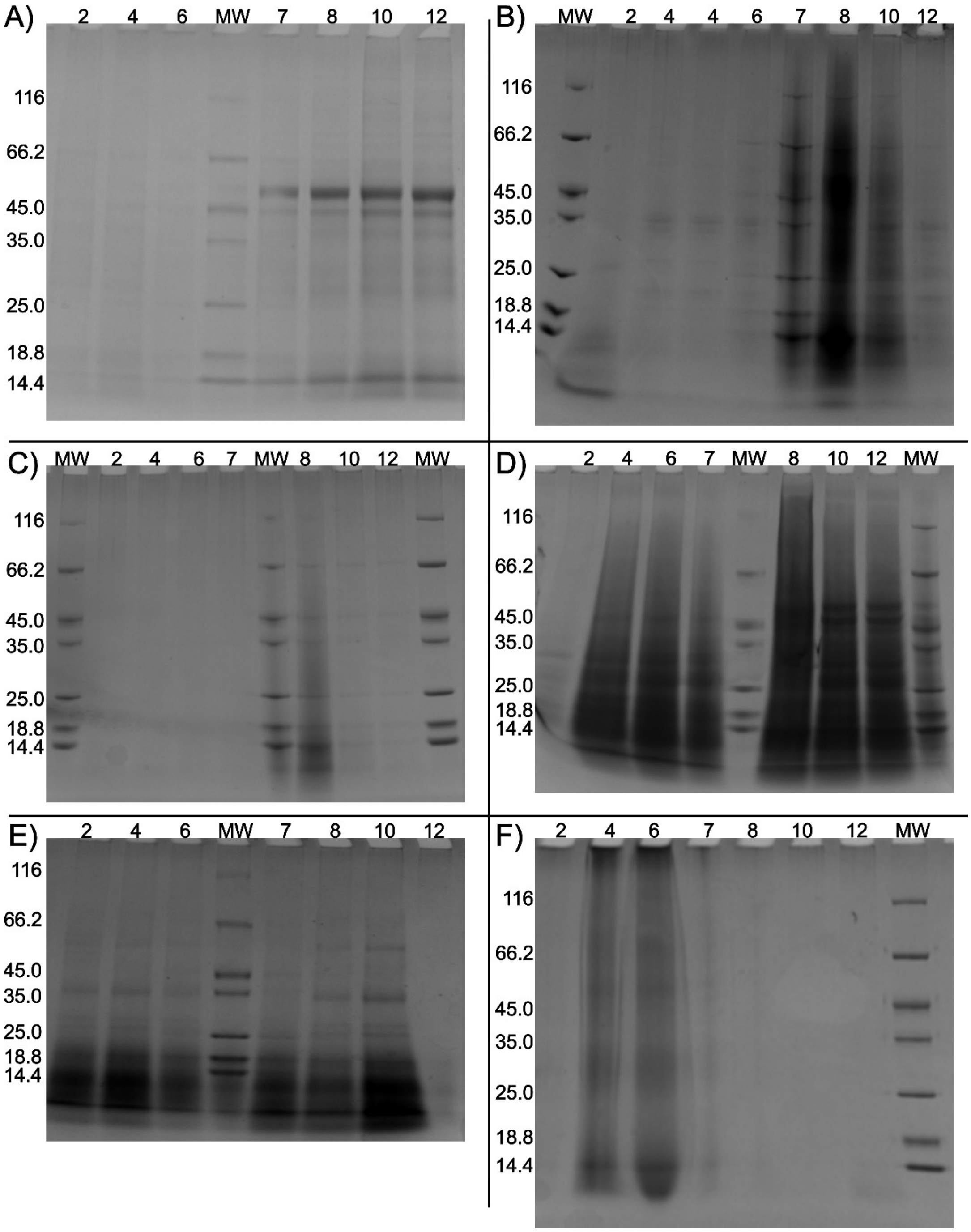


**Figure S1:** Visualization of **s**oluble protein at different pH (from solubility test) of the different clover grass protein (GCP) prototypes by SDS-PAGE. A: First stage membrane filtration permeate (MF). B: Acid precipitate (AP). C: Heat precipitate (HP). D: Second stage diafiltration concentrate (DC). E: DC hydrolysate (DCH). F: Heat and acid precipitate (HAP). "MW" refers to the molecular weight marker with MW of marker bands indicated. Above each lane, the pH of the sample loaded (equal volume loading) is annotated.

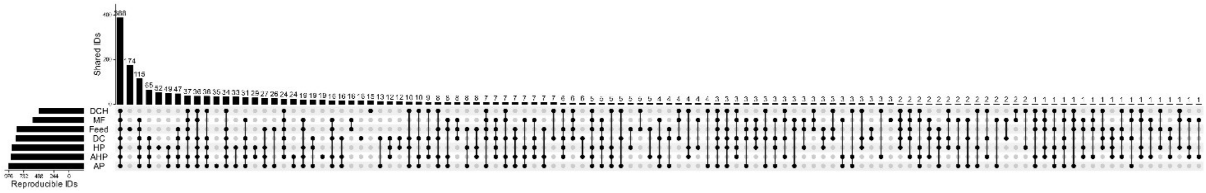

**Figure S2:** Full UpSet Plot from Fig. 4A. UpSet plot showing the number and overlap of reproducible (quantified in at least two of the replicates) protein IDs across the feed and GCP prototypes.

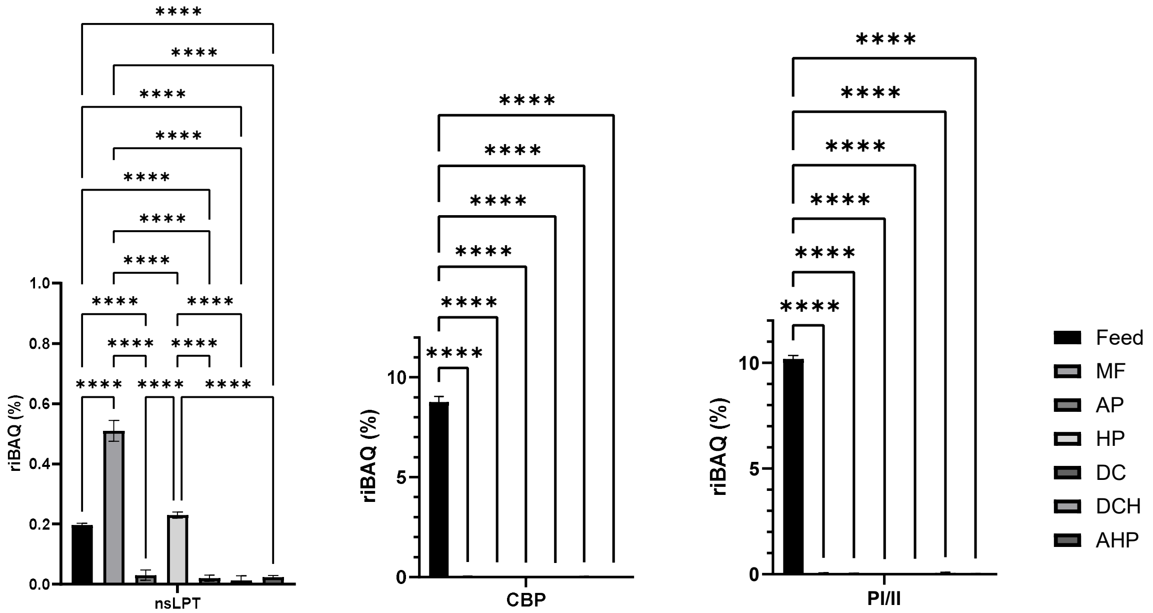


**Figure S3:** Differential abundance (by riBAQ) for non-specific lipid transfer proteins (nsLTPs), chorophyll-binding proteins (CBPs), and photosystem I and II proteins (PI/II) across all samples. Total abundance (sum of all subunits and isoforms) across the initial feed stream and all CGP prototypes for nsLTPs (left), CBPs (middle), and PI/II proteins (right). Abundance is indicated as ΣriBAQ (mean +/- SD) based on triplicate analysis for each sample. Statistical analysis is performed as one-way ANOVA with significance level (from adjusted p-values) indicated by "*" ($p \leq 0.05$), "**" ($p \leq 0.01$), "***" ($p \leq 0.001$), and "****" ($p \leq 0.0001$). Non-significant differences ($p > 0.05$), are not shown for simplicity.
